# Genetic interaction between *GL15* and *FDL1* modulates juvenile cuticle deposition and leaf permeability in maize

**DOI:** 10.1101/2024.10.21.619455

**Authors:** Giulia Castorina, Frédéric Domergue, Gabriella Consonni

## Abstract

The plant cuticle is a hydrophobic layer that serves as the primary barrier between plant surfaces and the external environment, limiting water loss and protecting against various stresses. This study investigates the genetic interaction between the maize *ZmFDL1* and *ZmGL15* genes and their role in modulating juvenile cuticle deposition and leaf permeability. The research was undertaken to understand how these genes influence cuticle composition and structure, which are crucial for plant protection against environmental stresses such as drought. We analysed the chemical composition of cutin and waxes, as well as cuticle-dependent leaf permeability. Our findings reveal that ZmFDL1 and ZmGL15 individually and synergically regulate the deposition of cuticular components, with significant impacts on leaf permeability. The *gl15-S* mutant exhibited an adult-like cuticle with higher cutin and lower wax contents, leading to reduced leaf permeability and improved water retention under drought conditions. These results highlight the importance of cutin in forming an effective water barrier and suggest that modulating cuticle composition with enhanced cutin content could be a strategy for improving drought tolerance in crops. This study provides new insights into the genetic regulation of cuticle biosynthesis and its relevance for plant adaptation to water scarcity.

**Highlight:**

- ZmFDL1 and ZmGL15 interact to modulate cuticle composition and reduce leaf water loss.
- *ZmFDL1* and *ZmGL15* genes regulate juvenile cuticle deposition in maize.
- The *gl15-S* mutant shows an adult-like cuticle with higher cutin and lower wax contents.
- The *gl15-S* mutation is epistatic to *fdl1-1*, partially rescuing the defective *fdl1* cuticle phenotype.
- The cuticle composition in *gl15-S* mutants leads to reduced leaf permeability and improved water retention under drought conditions.
- Cuticular layer with enhanced cutin content could improve drought tolerance in crops.

## INTRODUCTION

The plant cuticle is a hydrophobic layer produced and secreted by shoot epidermal cells, which forms the main barrier between the plant aerial surfaces and the external environment. The cuticle limits non-stomatal water loss and protects plants against numerous abiotic and biotic stresses, including drought, ultraviolet radiation, heat, and pest and pathogen invasion (Yeats and Rose, 2013, Xue *et al*., 2017; Ziv *et al*., 2018; Batsale *et al*., 2021).

Cutin and waxes are the two principal lipid components of the cuticle. Cutin, a cross-esterified polymer of glycerol backbones and hydroxylated- and epoxy-long chain (16 and 18 carbon atoms in length) fatty acids (Fich *et al*., 2016; Li-Beisson *et al*., 2013) forms an insoluble matrix, that is embedded by cuticular waxes. Cuticular waxes are a complex mixture of very-long-chain fatty acids (VLCFAs) of 20 or more carbon atoms in chain length, and their derivatives, which can include alcohols, aldehydes, alkanes, ketones and wax esters, as well as variable amounts of cyclic compounds, such as triterpenoids and phenylpropanoids (Bernard and Joubès, 2013; Lee and Suh, 2015). Cuticular waxes are also layered on top of the cutin matrix, forming epicuticular films or wax crystalloids (Bernard and Joubès, 2013; Yeats and Rose, 2013)

Cuticle composition and structure vary among plant species and organs of a single plant, as well as across developmental stages. In maize, cuticle composition and structure are among the traits that distinguish the leaves of the juvenile and adult phase. Depending on genotype, juvenile leaves are the first or six leaves produced by the plant. These leaves have a thin cuticle and are densely covered with epicuticular waxes crystals (Sylvester *et al*., 1990; Bongard-Pierce *et al*., 1996). Waxes of juvenile leaves mainly contain very-long-chain alcohols and aldehydes, and a lower proportion of alkanes and esters (Bianchi *et al*., 1978; Sturaro *et al*., 2005; Javelle *et al*., 2010; Loneman *et al*., 2017). In adult leaves, which have a thick cuticle and an amorphous wax layer on their surfaces, alkanes and alkyl esters are instead the principal wax components (Bianchi *et al*., 1984; Avato *et al*., 1990; Yang *et al*., 1993; Bourgault *et al*., 2020). Differences in cutin composition between juvenile and adult leaves have been poorly investigated and their impact on cuticle-dependent leaf permeability is still unknown.

Because of its protective functions, the cuticle is considered as a promising target for improving plant tolerance to environmental stresses, particularly drought. Cuticle composition and structure have been shown to influence cuticular permeability, and in turn drought tolerance, in various plant species. Genetic variants with defective cuticle composition have been frequently associated with increased cuticular permeability and reduced drought tolerance (Park *et al*., 2010; Zhou *et al*., 2013; Zhu and Xiong, 2013; Li *et al*., 2019), whereas overexpression of regulatory or structural genes of the cuticle biosynthetic pathways have been shown to cause increases in drought tolerance (Aharoni *et al*., 2004; Zhang *et al*., 2007; Kosma *et al*., 2009; Cui *et al*., 2016). Moreover, the expression of regulatory genes, which stimulates the activation of cuticle-biosynthesis-related genes, was shown to be promoted by drought (Seo *et al*., 2009; Seo *et al*., 2011; Lee and Suh, 2015; Bi *et al*., 2016).

Transcriptional regulators of cuticle biosynthesis, such as *ZmGLOSSY15* (*ZmGL15*), *ZmGLOSSY3* (*ZmGL3*), *ZmMYB94*/*FUSED LEAVES1* (*ZmFDL1*) and *ZmMYB84* have been detected in maize through mutant studies (Moose and Sisco, 1994; Liu *et al*., 2012, La Rocca *et al*., 2015; Liu *et al*., 2022) and, in the case of OUTER CELL LAYER1 (ZmOCL1), through the analysis of transcription patterns (Javelle *et al*., 2010). In previous studies, we have characterized the ZmMYB94/FUSED LEAVES1 (ZmFDL1) transcription factor as a key regulator of cuticle deposition in maize seedlings (La Rocca *et al*., 2015; Castorina *et al*., 2020). Seedlings lacking ZmFDL1 show a decrease in epicuticular wax load, mainly due to a reduction in primary very-long-chain alcohols, as well as in the cutin content, which is mainly attributed to decreases in ω-hydroxy fatty acids and polyhydroxy-fatty acids. These cuticle defects impact seedling development as well as leaf water holding capacity (La Rocca *et al*., 2015; Castorina *et al*., 2020).

*ZmGLOSSY15* (*ZmGL15*), an APETALA-2 like gene, was shown to play a fundamental role in promoting epidermal cell traits that define juvenile leaf identity and suppress adult leaf identity traits (Moose and Sisco, 1994; Moose and Sisco, 1996). *ZmGL15* is controlled by *miR172* that, by down regulating *ZmGL15*, promotes the transition from juvenile to adult vegetative phase (Lauter *et al*., 2005). In *gl15* mutant plants, the dull appearance of wild-type juvenile leaves is replaced by the glossy phenotype which is peculiar of adult leaves.

Previously, we reported that *ZmFDL1* and *ZmGL15* show similar variations of their expression profiles in response to drought. Under water scarcity conditions, both *ZmFDL1* and *ZmGL15* expression patterns exhibited an anticipation of the maximum transcript level peak compared to well-watered plants (Castorina *et al*., 2020). In the present study, we provide a comprehensive picture of the role of the *ZmGL15* gene in controlling chemical composition, structure, and function of the juvenile maize cuticle. We show that these cuticle-related traits are controlled by a genetic interaction between *ZmFDL1* and *ZmGL15*. In maize seedlings, cuticle features of the juvenile leaves, as well as the extent of cuticle-mediated leaf permeability, depend on both the *ZmFDL1* and *ZmGL15* individual genetic constitution. Moreover, data obtained from the chemical analysis of wild-type, single and double mutant genotypes have allowed to identify some correlations between variation in cuticle components and changes in cuticle-mediated leaf permeability. Overall, these results provide new insights into the identification of the cuticle components that provide better protection against water loss. This information may be useful to generate new strategies for improving plant response to drought stress as well as plant adaptation to water scarcity conditions.

## MATERIALS AND METHODS

### Plant Materials and Growth Conditions

The previously described maize (*Zea mays*) *fdl1-1* mutant (Castorina *et al*., 2020) used in this work was originally identified in the selfed progeny of a maize line crossed as female to an En/Spm line (La Rocca *et al*., 2015). The maize *gl15* mutant (*gl15-Sprague*, *gl15-S*) seeds were obtained from the Maize Genetics COOP Stock Center (catalogue no. 917E; maizecoop.cropsci.uiuc.edu) (Portwood *et al*., 2019). The other *gl15* mutant alleles (*gl15-Hayes, gl15-H* and *gl15-Lambert, gl15-L*) were kindly provided by Stephen Moose and were previously detailed (Moose and Sisco, 1994; Moose and Sisco, 1996; Lauter *et al*., 2005). To generate double-homozygous *fdl1-1 gl15-S* plants, single-homozygous *fdl1-1* and *gl15-S* plants were crossed. In all the experiments performed, homozygous mutants and their wild-type control plants were from the same segregating family. Plants were grown in a growth chamber with controlled temperature (25°C night/30°Cday) under a long-day photoperiod (16 h light/8 h dark) and with photon fluence of 270 µmol m^-2^ s^-1^.

To measure leaf water content under stress, maize seedlings were germinated and grown in soil under well-watered conditions until 17 DAS (time point 0). Then the drought was imposition by withholding irrigation.

### Epidermal Cell Density and Stomatal index

Epidermal cell density and stomatal index were measured using the surface imprint method on the abaxial and adaxial sides of the mid portion of fully expanded leaves, as previously described (Castorina *et al*., 2018). We analysed the mid portion of the second and third fully expanded leaves of wild-type and gl15-S mutant plants. The stomatal index was determined as (number of stomata/[number of epidermal cells + number of stomata]) × 100.

### Toluidine Blue Staining of Maize Epidermal Peels

Epidermal peels were sampled by hand from the mid portion of the second, third and fourth leaves of wild-type, *gl15-S*, *fdl1-1* and *fdl1-1 gl15-S* plants. The staining was performed as in the protocol described in Dudley and Poethig (1993) but slightly modified as follows: the fresh tissues were soaked in tap water and then stained for 5 minutes in a 1:1 aqueous solution of 0.05% toluidine blue O and 0.1M phosphate buffer at pH 6. The stained peels were washed for 1 minute in tap water and photographed using a light microscope (Olympus BX50) at 40x magnification. Juvenile and adult epidermal cells stained violet/pink or aqueous/turquoise, respectively.

### Cuticular Cutin and Wax Analysis

Cuticle composition was analysed in the first, second and third mature leaves of homozygous *gl15-S* and its wild-type control seedlings. The cuticular cutin and wax composition was also analysed in the second leaves of double-homozygous *fdl1-1 gl15-S*, *fdl1-1* and single-homozygous *gl15-S* and wild-type plants, respectively. Cutin and epicuticular waxes were extracted and identified by the combination of gas chromatography on-column injection and gas chromatography-mass spectrometry performed according to previously described methods (Domergue *et al*., 2010; Bourdenx *et al*., 2011).

### Chlorophyll Leaching Assay, Water Loss and Leaf Relative Water Content

The chlorophyll leaching assay was performed on mutant and wild-type fully expanded leaves. Leaf samples were immersed in 80 % ethanol and chlorophyll released was quantified in samples taken from the solution by measuring the absorbance at 647 and 664 nm. The concentration of total chlorophyll was calculated using the equation described by Lolle *et al*., (1997). Biological replicates consisted in leaves taken from independent plants.

To determine the rate of the leaf water loss, leaves were detached and weighed immediately. Sample weight was then estimated at time intervals and water loss was calculated as the percentage of fresh weight based on the initial weight.

To monitor the leaf moisture content under drought stress, the Relative Water Content (RWC) (Smart and Bingham, 1974) was measured in wild-type control and *gl15-S* mutant plants. The second and third leaf RWC was measured at 0, 24 and 48 hours after withholding irrigation taking a leaf blade disc of 0.5 cm^2^ from the median portion of the leaf. Fresh weight (FW) was weighed immediately after samples collection. The total weight (TW) was obtain soaking the tissue for 24 hours in distilled water. Then, the discs were fully dried and eventually, the dray weight (DW). The leaf RWC was calculates with following formula: RWC = ([FW-DW]/[TW-DW]) x 100. For each treatment, the leaf RWC was measured a minimum of four independent plants per genotype.

### SEM Analyses of Epicuticular Waxes

For the analysis of epicuticular waxes at the scanning electron microscope (SEM), samples were air-dried and processed according to Castorina *et al*. (2018; 2023). Leaf pieces from the mid portion were fixed using double-sided tape to a carbon base on aluminium stubs (ø 12.5 mm; 3.2 x 8 mm pin) and gold metalized using a Scancoat Six Sputter Coater (Edwards). Micrographs of the abaxial and adaxial leaf surfaces were acquired with the SEM-EDS JSM-IT500 LV electron microscope (JEOL Spa).

### RNA extraction and Gene Expression Analysis

Total RNA was extracted with the TRIzol Reagent (Life Technologies), suspended in RNase-free milliQ dH2O and treated with RQ1 RNase-Free DNase (Promega). RNA concentration was measured with the NanoDrop® ND-1000 spectrophotometer (Thermo Scientific). First-strand cDNA was synthetized using the High-Capacity cDNA Reverse Transcription Kit (Thermo Fisher Scientific). Quantitative Real-Time PCR (RT-qPCR) was performed as previously described (Castorina *et al*., 2020). The gene-specific primers are listed in Supplementary Table S1.

For the expression enrichment test, B73 wild-type seedlings were grown on soil and at 10 DAS, the second leaf was sampled. The epidermis was manually peeled off, with the help of tweezers, and immediately frozen in liquid nitrogen. From the remaining portion of the leaf, with the help of a spatula the green tissues, consisting mostly of mesophyll, was sampled. Also, the whole leaf was sampled. Four biological replicates were sampled per thesis and each replicate consists of leaves tissues from three independent plant.

### Statistical analysis

All data were statistically analysed. Student’s t test (ns = not significant, \**P* < 0.05, \*\**P* < 0.01, *** *P* < 0.001 and *** *P* < 0.0001) or two-way analysis of variance (ANOVA) (*P* < 0.05), combined with Post-hoc Tukey tests, was conducted using the statistical packages Prism GraphPad 10.

### Accession Numbers

The gene sequences from this article can be found in the Maize Genetics and Genomics Database (MaizeGDB) or GenBank/EMBL databases under the following accession numbers: Zm00001eb328280 (*ZmFDL1/MYB94*), Zm00001eb387280 (*ZmGL15*), Zm00001eb042160 (*ZmHTH1*), Zm00001eb311010 (*ZmONI3*), Zm00001eb195850 (*ZmGL3*), Zm00001eb246270 (*ZmGL8*), Zm00001eb190120 (*ZmGL4*), Zm00001eb018600 (*ZmKCS39*), Zm00001eb296230 (*ZmKCS16*), Zm00001eb176110 (*ZmCER4*), Zm00001eb247450 (*ZmCER1*), Zm00001eb156680 (*ZmWSD1*), Zm00001eb313510 (*ZmGL1*), Zm00001eb071110 (*ZmGL2*), Zm00001eb122470 (*ZmGL13*), Zm00001eb087050 (*ZmGL14*), Zm00001eb385900 (*ZmEF1*α).

## RESULTS

### Spatiotemporal expression analysis of *ZmFDL1* and *ZmGL15* genes

We have previously reported that *ZmFDL1* and *ZmGL15* transcription factors are both expressed in maize seedling leaves and are drought responsive (Castorina *et al*., 2020). To further define their pattern of expression, quantitative Real-Time PCR (RT-qPCR) was performed (Fig. 1). In the second leaf of wild-type seedlings at 7, 9 and 11 days after sowing (DAS), expression patterns were similar and showed a progressive increase in the transcript levels, but the expression levels of *ZmGL15* were much higher compared to *ZmFDL1* (Fig. 1A).

**Figure 1.**
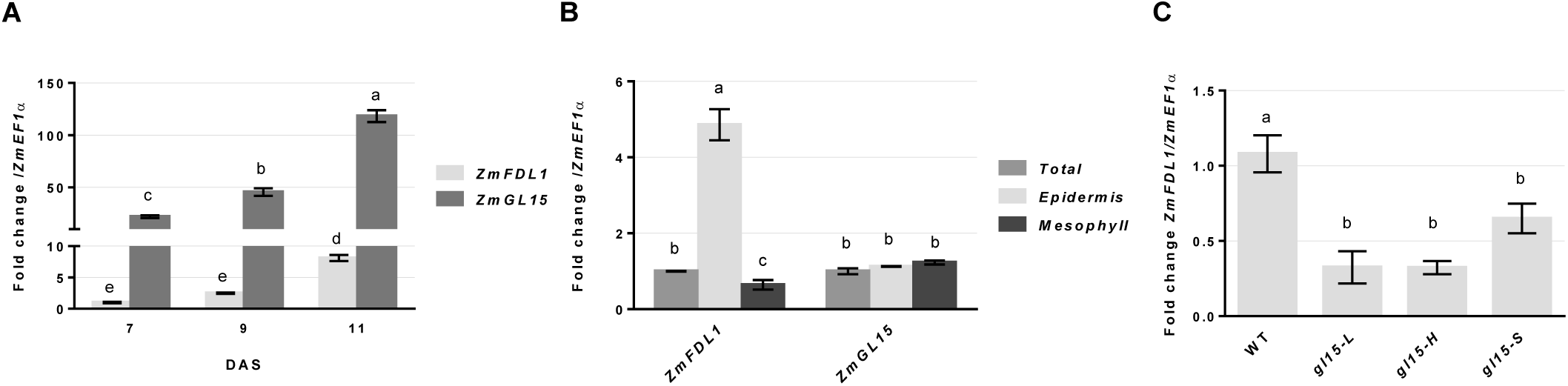
gene expression analysis of *ZmGL15* and *ZmFDL1*. Transcript level quantification of *ZmGL15* and *ZmFDL1* genes performed, by RT-qPCR, (A) in the second leaf of wild-type plants at 7, 9 and 11 DAS and (B) in the whole leaf (Total), mesophyll and epidermal tissues from the second leaves of 10 DAS wild-type plants. (C) Gene expression level of *ZmFDL1* in homozygous *gl15* mutants and wild-type (WT) control seedlings at the coleoptile developmental stage. Values represent the mean fold change variations of a minimum of four biological replicates. Error bars are ±SE. Different letters denote significant differences assessed by two-way ANOVA (*P*<0.05).

We also performed an expression enrichment test in which different tissues from the mature second leaf of 10 DAS seedlings were sampled. We observed that the *ZmFDL1* transcript level was significantly higher in the epidermis as compared to the mesophyll-enriched and whole leaf samples (Fig. 1B), indicating that *ZmFDL1* is specifically expressed in the epidermal layer. This observation is consistent with previous findings (Liu *et al*., 2021) and is coherent with *ZmFDL1* specific involvement in the control of cuticle deposition, a process which occurs exclusively in epidermal cells. Moreover, this result confirmed the robustness of this approach. No differences were instead detected among whole leaf, mesophyll and epidermal tissue samples for the transcript levels of *ZmGL15* (Fig. 1B), suggesting that this gene does not exhibit preferential expression in a specific leaf tissue layer.

The ZmFDL1 transcription factor is a key regulator of cuticle deposition in juvenile vegetative tissues. Lack of ZmFDL1 activity affects the expression of several cuticular wax genes involved in different modules of the biosynthetic network (Castorina *et al*., 2020). Among the cuticle-related genes which were shown to be differentially expressed in *fdl1-1* mutant seedlings, 14 were also analysed for their tissue-dependent expression specificity. The expression level of most of these genes was found to be much higher in the samples enriched with the epidermal tissue (Supplementary Fig. S1), as already shown in Arabidopsis stems (Suh *et al*., 2005).

We previously showed that the expression of *ZmGL15* was lower in the *fdl1-1* mutant compared to wild type (Castorina *et al*., 2020). In this work, a significant downregulation of *ZmFDL1* was observed in three homozygous *gl15* genotypes that carried different *ZmGL15* mutant alleles, i.e., *gl15-L*, *gl15-H* and *gl15-S* (Fig. 1C). This indicated that *ZmGL15* promotes the expression of *ZmFDL1,* and that the two transcription factors reciprocally stimulate their expression.

### Analysis of cuticle-related leaf traits in *gl15-S* mutant plants

Mutations in the *ZmGL15* gene were shown to cause the coordinate precocious expression of adult epidermal cell features (Moose and Sisco 1994). Accordingly, we observed the precocious presence of the glossy phenotype in *gl15-S* mutant seedlings, which was evident starting from the third leaf (Fig. 2A). Third and fourth leaves of *gl15-S* seedlings appeared glossy, or shiny green, in contrast with the dull appearance of the corresponding wild-type leaves (Fig. 2A). In addition, when leaves were sprayed with water, water droplets adhered to the *gl15-S* mutant leaf surfaces whereas, the water slipped away on wild-type leaves (Fig. 2A, insets). More precisely, the glossy phenotype showed a tip-base gradient in the third leaf of the *gl15-S* mutant. The base-to-middle portion of the leaf displayed an obvious glossy phenotype while the middle-to-tip leaf portion showed a decrease in the glossy phenotype intensity and the leaf tip appeared dull. This suggests that this leaf is undergoing the transition from juvenile to adult phase.

**Figure 2.**
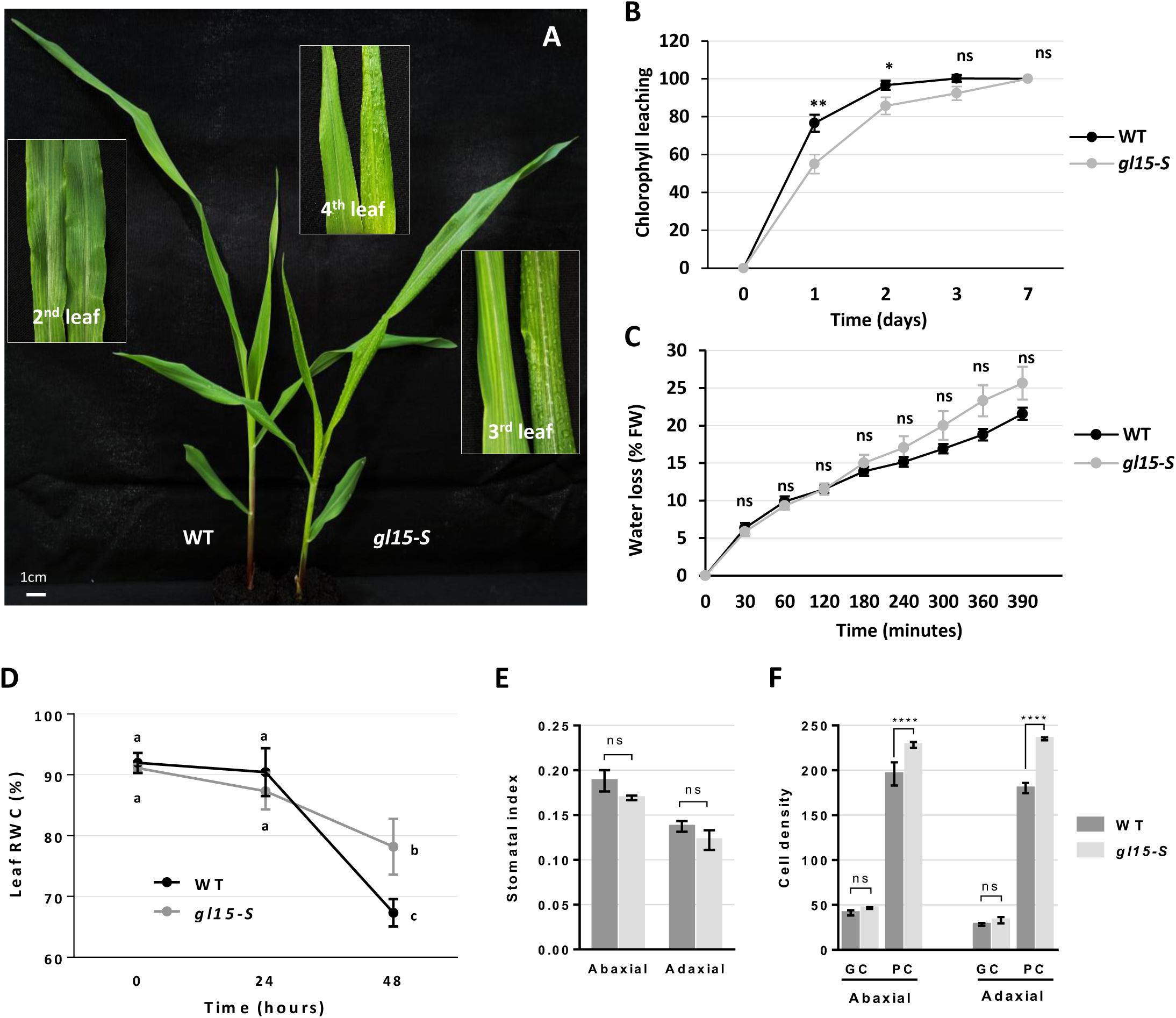
Cuticle-dependent phenotype in *gl15-S* and wild-type plants. (A) Representative image of wild-type (WT) and homozygous *gl15-S* 15-day-old seedlings. The insets represent a magnification of wild-type (left) and *gl15-S* (right) second, third and fourth leaves. The glossy phenotype and the ability of the leaf surface to retain water drops were visible starting from the third leaf of *gl15-S* mutant plants. (B) The chlorophyll leaching assay and (C) leaf evapotranspiration, expressed as the percentage of water loss of fresh weight (FW), were performed on the second fully expanded leaf of wild-type (WT) and homozygous *gl15-S* 15-day-old plants. Values represent the mean of 10 biological replicates per genotype. Error bars are ±SE. Differences were assessed by the Student’s T-test (* *P*<0.05; ** *P*<0.01; ns=not significant). (D) The relative water content (RWC) was measured in the second leaf of wild-type (WT) and *gl15-S* plants subjected to 24 and 48 hours of water scarcity imposed by withholding irrigation. Values represent the mean of a minimum of four biological replicates. Error bars are ±SE. Different letters denote significant differences assessed by two-way ANOVA (*P*<0.05). (E) Stomatal index and (F) the density per 1 mm^2^ of guard cells (GC) and pavement cells (PC) were measured in both the abaxial and adaxial sides of the second leaf of wild-type (WT) and *gl15-S* plants. Values are the mean ± SE and differences were evaluated by Student’s T-test.

In addition, we analysed the epidermal traits of the adaxial and abaxial sides of second and third leaves in the middle portion of the wild-type and *gl15-S* mutant plants. Juvenile epidermal traits, *i.e.* cells with wavy walls and a pink coloration of epidermal peels following toluidine blue-staining, were observed in both leaves of wild-type plants (Supplementary Fig. S2 A-D; I-L) and in the second leaf of *gl15-S* mutant plants (Supplementary Fig. S2 E-H). In contrast, the *gl15-S* mutant plants expressed both juvenile and adult epidermal cell traits, the latter including longer epidermal cells with highly crenulated lateral walls (Supplementary Fig. S2M, O) and aqua/turquoise-coloured areas in the toluidine blue-stained epidermal peels (Supplementary Fig. S2N, O). The results confirmed that the second leaf of the *gl15-S* mutant was juvenile while the third leaf was in transition, showing epidermal cells with traits of both phases.

To further investigate the impact of the *gl15-S* mutation on leaf cuticle, leaf permeability was first assessed using the chlorophyll leaching assay. The first (Fig. 2B), second (Supplementary Fig. S3A) and third (Supplementary Fig. S3C) leaves of *gl15-S* seedlings released chlorophyll slower as compared to wild-type control siblings indicating a lower cuticle-dependent leaf permeability, whereas no statistical differences were observed for the fourth leaf (Supplementary Fig. S3E). Then, a water loss time course experiment was performed on wild-type and *gl15-S* leaves by estimating the rate of leaf weight loss in relation to the initial fresh weight. In all examined leaves, the resulting *gl15-S* mutant profiles were comparable to that of wild-type leaves (Fig. 2C; Supplementary Fig. S3B, D, F), suggesting that non-stomatal leaf evapotranspiration was not altered by the *gl15-S* mutation.

In contrast, leaf relative water content (RWC) analysis of plants grown under well-watered conditions or exposed to drought stress, which was applied by withholding irrigation, revealed differences between wild-types and *gl15-S* mutant plants (Fig. 2D; Supplementary Fig. S3G). After 48 hours of water scarcity, the leaf RWC measured in the second (Fig. 2D) and third (Supplementary Fig. S3G) leaves was higher in *gl15-S* mutants as compared to wild type. We also investigated if the increased water-holding capacity of the *gl15-S* leaves was influenced by stomatal traits. To this aim, the stomatal index (Fig. 2E; Supplementary Fig. S3H) and stomatal density (Fig. 2F; Supplementary Fig. S3I) were measured. However, no statistical differences were detected, suggesting that cuticular traits may be responsible for the different leaf RWC phenotypes observed.

Finally, we examined whether the abundance and shape of epicuticular wax crystals were affected in the *gl15-S* mutant using scanning electron microscopy (SEM). These analyses were conducted using the mid portion of the third leaf of wild-type and *gl15-S* mutant plants (Fig. 3). Fewer wax crystals were observed on both abaxial (Fig. 3F, J) and adaxial (Fig. 3H, L) surfaces of the *gl15-S* mutant glossy leaves relative to the surfaces of wild-type dull leaves (Fig. 3E, I and G, K), and these crystalloids appeared much smaller, flattened and embedded in the underneath cuticle layer (Fig. 3F, J, H, L). A similar reduction in size and distribution of the epicuticular wax crystals was previously observed on glossy leaves of maize mutants in cuticle-related genes (Beattie and Marcell, 2002; Li *et al*., 2019; Zheng *et al*., 2019; Liu *et al*., 2023).

**Figure 3.**
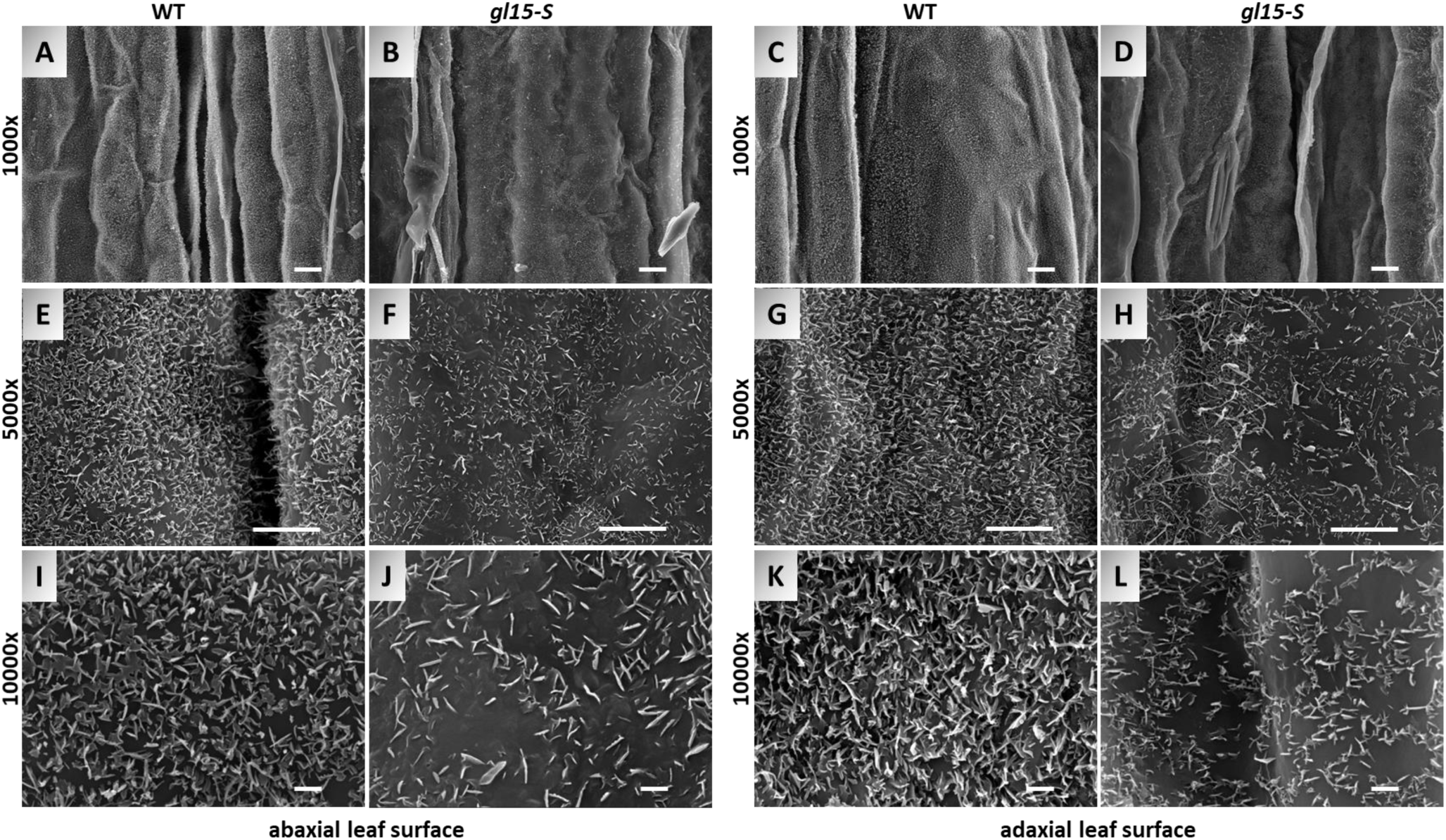
Distribution of cuticular waxes on the leaf surfaces of the *gl15-*S mutant. SEM micrograph images of both abaxial and adaxial surface of the third fully expanded leaf in wild type (WT) control plant (A, E, I, C, G, K) and single homozygous *gl15-S* (B, F, J, D, H, L) mutant have been acquired at 1000x, 5000x and 10000x magnification. Scale bars correspond to 10 µm (A-D), 5 µm (E-H) and 1 µm (I-L).

Overall, these results confirm that *gl15-S* mutation causes alterations in both structure and function of the juvenile cuticle. Interestingly, the observed leaf permeability was lower and, under drought stress, the leaf RWC was higher in leaves of the *gl15-S* mutant as compared to wild-type control siblings, thus suggesting that cuticle composition of the *gl15-S* mutant could be more efficient in preventing water loss in drought conditions.

### ZmGL15 regulates both cutin and wax deposition

To examine the effects of the *gl15-S* mutation on the cuticle composition of juvenile leaves, cutin and waxes were extracted from the first, second and third leaves of *gl15-S* mutant and wild-type plants and their analysis was performed by gas chromatography (Fig. 4) (Domergue *et al*., 2010; Bourdenx *et al*., 2011).

**Figure 4.**
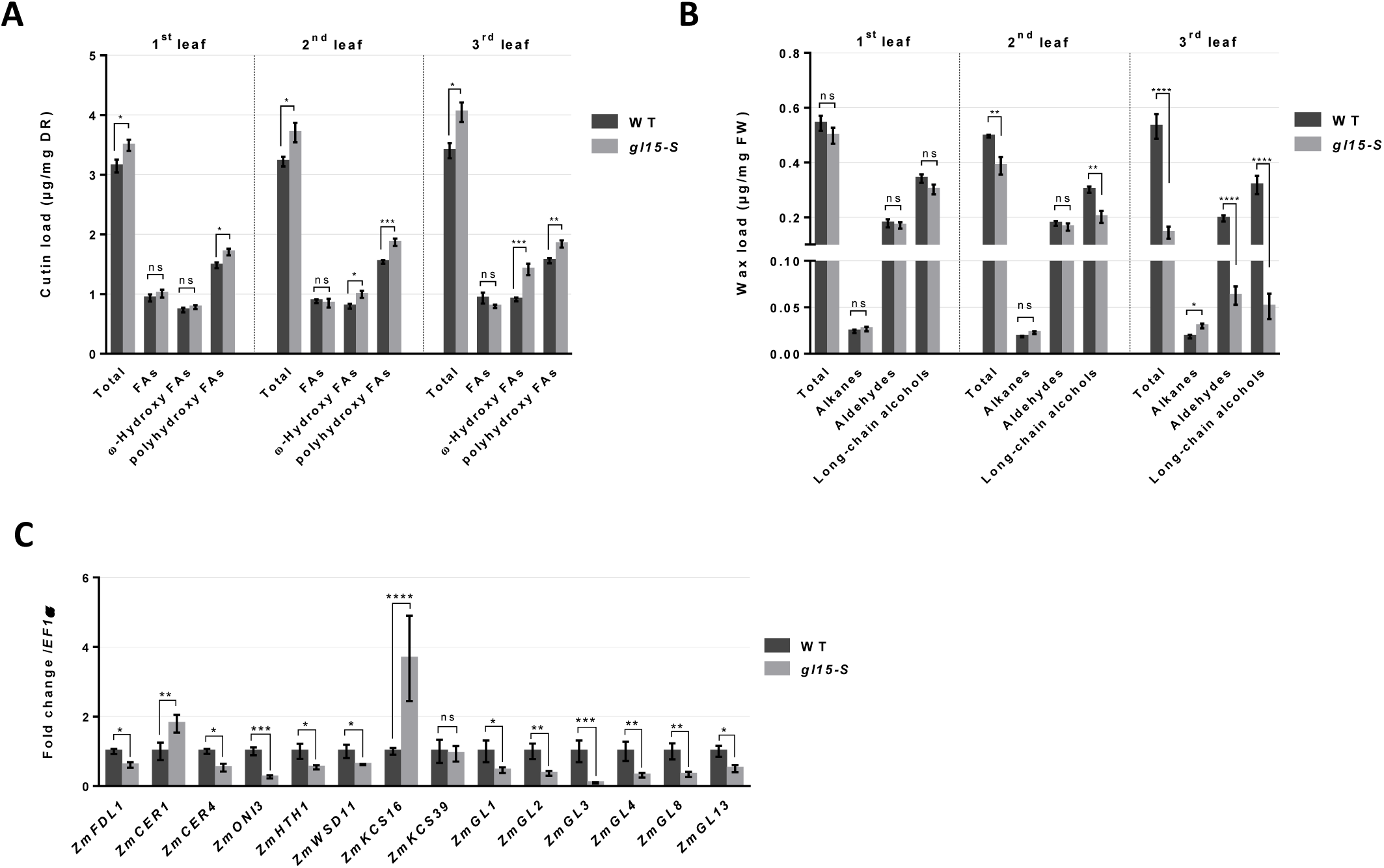
Cuticle composition in *gl15-S* plants. (A) Cutin aliphatic and (B) cuticular wax content of the first, second and third fully expanded leaf from 21-day-old *gl15-S* and wild-type (WT) plants. Values represent the mean of six biological replicates per genotype ± SE. (C) Gene expression analysis of cuticle-related genes. The transcript level of *ZmFDL1, ZmCER1, ZmCER4, ZmONI3, ZmHTH1, ZmWSD11, ZmKCS16, ZmKCS39, ZmGL1, ZmGL2, ZmGL3, ZmGL4, ZmGL8* and *ZmGL13* genes were analysed by RT-qPCR in the third leaf of homozygous *gl15-S* mutants and wild-type (WT) control plants. Values represent the mean fold change of four independent biological replicates. Error bars are ±SE. Comparison was made between genotypes and significant differences were assessed by Student’s T-test (* *P*<0.05; ** *P*<0.01; *** *P*<0.001; **** *P*<0.0001; ns=not significant).

Considering total cutin monomers amounts, a higher cutin load was observed in all three examined leaves of the *gl15-S* mutants as compared with wild-type leaves (Fig. 4A). While no differences were detected in the content of fatty acids (FAs) (Fig. 4A; Supplementary Fig. S4C), an increase in ω-hydroxy fatty acids (FAs) as well as in polyhydroxy FAs (Fig. 4A) was observed in *gl15-S* mutant compared to wild-type samples. The increase in ω-hydroxy FAs was mainly associated with the progressive increased accumulation of the 18:1 ω-hydroxy FA isomer (18:1ωOH; Supplementary Fig. S4D), while an increase in the content of several polyhydroxy FAs was observed in the three mutant leaves (Supplementary Fig. S4E). Also, considering relative abundance of the total cutin content, a progressive decrease in FAs and an increase in ω-hydroxy FAs were observed in *gl15-S* mutant seedling leaves (Supplementary Fig. S4A). For example, in the third leaf, the cutin of *gl15-S* mutant contained 19,7% of FAs and a 34.7% of ω-hydroxy FAs compared to the 27.1% and 26.9% in the wild-type, respectively (Supplementary Fig. S4A).

Concerning cuticular waxes, no differences were detected in the first leaf (Fig. 4B), but the *gl15-S* mutation caused a slight and strong reduction in the total wax load in the second and third leaves, respectively. Among wax metabolite classes, a decrease in alcohols in the second leaf, and in both aldehydes and alcohols in the third leaf (Fig. 4B) were observed. The analyses of individual wax metabolites revealed a statistically significant reduction in the C32 n-aldehyde content, which was observed only in the third leaf (ALD32; Supplementary Fig. S4G), and in the content of C32 primary alcohols (C32OH; Supplementary Fig. S4H), which was detected in both the second and third leaves. This change was greater in the third leaf, in which a reduction in very-long-chain C30 primary alcohols (C30OH; Supplementary Fig. S4H) was also observed. A slight increase in alkanes content was in contrast observed in the third leaves (Fig. 4B), which was due to the higher accumulation of C31 and C33 alkanes (ALK31 and ALK33; Supplementary; Fig. S4F). Considering relative abundance in total waxes, alkane and aldehyde increase, while long-chain alcohols decrease from the first to the third leaf of *gl15-S* mutant compared to wild-type seedlings (Supplementary Fig. S4B). In the third leaves, alkanes accounted for 23.8% and 3.5% of the total wax loads, aldehydes for 43.3% and 37.3%, and alcohols for 32.9% and 59.2% in the *gl15-S* mutant and wild-type controls, respectively (Supplementary Fig. S4B).

Variations in cuticle composition indicate the involvement of ZmGL15 in the control of cuticle-related gene expression. This was confirmed by analysing the expression levels of 14 cuticle-related genes, including *ZmFDL1,* in the third leaf of both wild-types and *gl15-S* mutants (Fig. 4C). Twelve out of fourteen genes showed a significant lower level of expression, while only two, *ZmCER1* and *ZmKCS16*, were upregulated in the *gl15-S* mutant. Only the expression levels of *ZmKCS39* did not significantly differ. The transcript levels of the MYB regulatory genes *ZmFDL1* (La Rocca *et al*., 2015) and *ZmGL3* (Liu *et al*., 2012), and of the *ZmGL13* gene encoding an ABC transporter (Li *et al*., 2013), were down regulated (Fig. 4C). Similarly, the expression of *ZmGL1* (Sturaro *et al*., 2005), *ZmGL2* (Tacke *et al*., 1995; Alexander *et al*., 2020), *ZmGL4* (Liu *et al*., 2009) and *ZmGL8* (Dietrich *et al*., 2005) genes, which are all involved in the FAs metabolism, seems to be stimulated by ZmGL15, since they were down regulated in the *gl15-S* mutant leaves (Fig. 4C). The cutin-related *ZmONI3* and *ZmHTH1* genes, which are both presumably involved in the synthesis of α, ω-dicarboxylic acids (DCAs), were also downregulated (Fig. 4C). As to genes related to wax biosynthesis, *ZmCER4* and *ZmWSD11*, putatively involved in the synthesis of long chain primary alcohols and of long chain wax ester respectively, were both down regulated. In agreement with our wax analyses, the *ZmCER1* gene, which is involved in the production of long-chain alkanes, was upregulated (Fig. 4C).

### Analysis of cuticle-dependent developmental defects shows that *gl15-S* is epistatic to *fdl1-1*

To examine the genetic relationship between *ZmGL15* and *ZmFDL1*, F2 progenies were produced from selfing heterozygous *gl15-S* /+ *fdl1-1* /+ F1 plants. A first analysis was performed during seedling development through visual scoring of the F2 plant phenotypes (Supplementary Fig. S5A). Four phenotypic classes were detected, comprising wild-type, two distinct single-homozygous mutant classes, *i.e.* gl15 and fdl1 respectively, and double-homozygous *gl15-S fdl1-1* mutant plants exhibiting both mutant traits (Supplementary Fig. S5A). The gl15 phenotype observed was as described in Figure 2, while *fdl1-1* mutant plants were recognizable for their impaired growth, caused by the presence of fusions between the coleoptile and first leaf, and between leaves (La Rocca *et al*., 2015).

Concerning seedling height, no differences were detected between wild-types and *gl15-S* mutants, at both 7 and 11 DAS (Fig. 5A). A statistically significant reduction in the elongation was instead observed for both *fdl1-1* and *fdl1-1 gl15-S* mutants as compared to both wild types and *gl15-S* mutants (Fig. 5A; Supplementary Fig. S5A,). This reduction was greater in *fdl1-1* than in *fdl1-1 gl15-S* homozygotes (Fig. 5A). An in-depth analysis of the *fdl1-1* mutants disclosed a variability in the expressivity of the phenotype that was consistent with previous observations (La Rocca *et al*., 2015). Specifically, we detected four mutant categories referred to as mild, intermediate, strong and severe phenotypes, respectively (Supplementary Fig. S5B). The latter category (13,8%) was only detected in the *fdl1-1* mutants (Supplementary Fig. S5C), and not found among the *fdl1-1 gl15-S* mutant plants (Supplementary Fig. S5D). In double mutants, we also observed instead a strong increase in the percentage of mild and intermediate fdl1 phenotypes: from 23.9% to 40.2% and from 25.5% to 44.7% *fdl1-1* and *fdl1-1 gl15-S* mutants, respectively. Consequently, the frequency of the strong phenotype decreased from 36.7% to 15.1% (Supplementary Fig. S5C, D).

**Figure 5.**
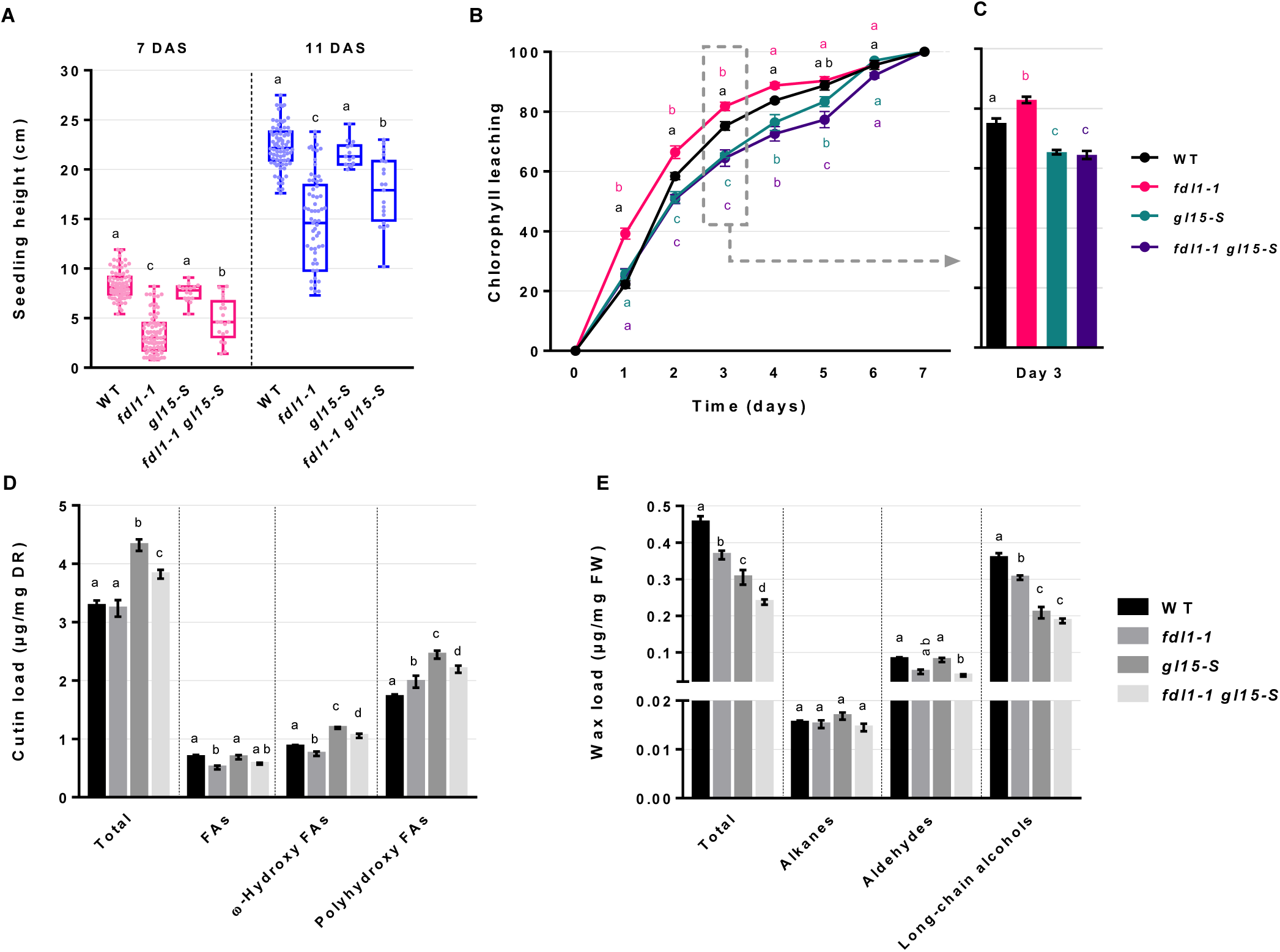
Effects of ZmGL15 and ZmFDL1 on cuticle-related traits. (A) Box plot of seedling height of wild-type (WT), single homozygous *fdl1-1*, *gl15-S* and double homozygous *fdl1-1 gl15-S* plants at 7 (red) and 11 (blue) DAS. Values represent the mean of minimum 15 biological replicates per genotype. (B) Chlorophyll leaching assay on the second fully expanded leaf of wild-type (WT), single homozygous *fdl1-1*, *gl15-S* and double homozygous *fdl1-1 gl15-S* plants. Values represent the mean of a minimum four biological replicates per genotype. Error bars are ±SE. (C) Cuticle-dependent leaf permeability at day 3 of the assay is shown in panel B. Error bars are ±SE. (D) Cutin aliphatic and (E) cuticular wax content of the second fully expanded leaf from 15-day-old plants. Values represent the mean of six biological replicates per genotype ± SE. Different letters denote statistically significant differences between genotypes assessed by two-way ANOVA (*P* < 0.05).

It is also well known that in *gl15* mutants, the precocious shift to the adult vegetative phase caused an anticipation of the flowering time measured as leaf number (Moose and Sisco, 1994; Moose and Sisco, 1996; Lauter *et al*., 2005). Accordingly, compared to wild types, a reduction in the total leaf number in the *gl15-S* mutant was observed at flowering (Supplementary Fig. S5E). The total leaf number was not affected in the *fdl1-1*, while the *fdl1-1 gl15-S* mutant exhibited a phenotype similar that of *gl15-S* (Supplementary Fig. S5E). The reduced leaf number suggested a precocious shift from the juvenile to the adult vegetative phase also in the *fdl1-1 gl15-S* mutant.

The analysis of the epidermal traits on the abaxial sides of second (Supplementary Fig. S5F-I), third (Supplementary Fig. S5J-M) and fourth (Supplementary Fig. S5N-Q) leaves in all four genotypes, corroborated this hypothesis. Toluidine blue-staining of epidermal peels revealed that wild-type (Supplementary Fig. S5F, J, N) and *fdl1-1* mutant (Supplementary Fig. S5G, K, O) leaves retain juvenile epidermal traits up to the fourth leaf since all stained pink. In contrast, the epidermis of *gl15-S* and *fdl1-1 gl15-S* mutant plants exhibited pink-stained epidermal cells and juvenile traits in the second leaves only (Supplementary Fig. S5H, I), while showing pink/turquoise-staining and turquoise-staining in the epidermal tissues of the third (Supplementary Fig. S5L, M) and fourth leaves (Supplementary Fig. S5P, Q), respectively.

The analysis of seedling developmental defects (Fig. 5A; Fig. S5C, D) indicated that the *gl15-S* mutant can partially rescue the fdl1 phenotype. Furthermore, analysis of flowering time (Fig. S5E) and epidermal traits (Fig. S5F-Q) unambiguously demonstrates that *gl15-S* is epistatic to *fdl1-1*.

Further analyses were conducted on the second leaf that was chosen based on the following observations: *i*) differences in cuticle permeability between *fdl1-1* and the wild-type control plants were more pronounced in the second leaf (Castorina *et al*., 2020); *ii*) *ZmDFL1* and *ZmGL15* exhibited a similar temporal expression pattern in the second leaf (Fig. 1); *iii*) the glossy phenotype appears from the third leaf onwards in *gl15-S* mutant (Fig. 2A) indicating a maintenance of juvenile epidermal traits in the second leaf (Supplementary Fig. S5E-H); *iv*) despite its dull appearance, the second leaf of the *gl15-S* mutant showed a reduced cuticle-dependent permeability, accompanied by alterations in the chemical composition of the cuticle (Fig. 4). Altogether these findings highlighted that both genes are involved in the control of juvenile cuticle deposition in this leaf.

The chlorophyll leaching assay was conducted on the four genotypes. The results showed that the second leaf of *fdl1-1* mutants released chlorophyll faster than wild type and *gl15-S* mutants (Fig. 5B, C). Interestingly, chlorophyll leaching was equivalent in the *gl15-S* and *fdl1-1 gl15-S* mutants (Fig. 5B, C).

### ZmGL15 and ZmFDL1 regulate juvenile cuticle deposition

To elucidate the nature of the cuticle alterations that allow the rescue of the *fdl1-1* leaf permeability defects in the double *fdl1-1 gl15-S* mutant, the chemical composition of the cutin and cuticular waxes were analysed on the second fully expanded leaves of wild-type, and *gl15-S*, *fdl1-1* and *fdl1-1 gl15-S* mutant plants (Fig. 5C, D; Supplementary Fig. S6).

The total cutin load did not statistically differ in *fdl1-1* mutant plants compared to wild type (Fig. 5D). Both *gl15-S* and the *fdl1-1 gl15-S* mutants were statistically different compare to wild type and *fdl1-1* mutant but *gl15-S* mutant showed the highest increase in cutin load (Fig. 5D).

No statistical differences were detected in the total fatty acids (FAs) content between genotypes, except for a decrease in *fdl1-1* plants, while an increase in polyhydroxy FAs was observed in all the mutant genotypes compared to the wild type (Fig. 5D). The ω-hydroxy FAs content showed a reduction in *fdl1-1* mutant and an increased load in the *gl15-S* and *fdl1-1 gl15-S* compared to control plants (Fig. 5D). Considering the relative abundance of the different cutin compound classes, differences between genotypes have been detected. We observed a decrease in FAs (15%) percentage in all mutant seedling leaves compared to wild type (Supplementary Fig. S6A). The relative abundance of polyhydroxy FAs was much higher in the *fdl1-1* mutant (61.1%) compared to the wild type (52.4%), while the *gl15-S* and *fdl1-1 gl15-S* mutants exhibited a similar intermediate relative abundance (56.5% and 57.3%, respectively) (Supplementary Fig. S6A). Instead, no statistical variation in the relative content of the ω-hydroxy FAs was detected except for the single-homozygous *fdl1-1* mutant which exhibit a lower percentage (23%) compared to the other genotypes (26.5%-27.5%) (Supplementary Fig. S6A).

The effect of the *gl15-S* recessive allele on *fdl1-1* was also detectable in single cutin metabolites (Supplementary Fig. S6A, C-E). The amount of FA isomers was statistically lower in *fdl1-1* plants compared to the wild type but in *fdl1-1 gl15-S* the levels were restored to those of the control plants (Supplementary Fig. S6C). Considering the ω-hydroxy FA isomers, except for C18:1 and C24:0, the amounts in the double-homozygous *fdl1-1 gl15-S* mutant were equal to wild type even when the amount was lower in the single-homozygous *fdl1-1* mutant (Supplementary Fig. S6D). In contrast we observed that the amount of the 18:1 ω-hydroxy FAs (18:1ωOH), the most abundant cutin isomer, was much higher in the double-homozygous *fdl1-1 gl15-*S mutant compared to both wild-type and *fdl1-1* plants (Supplementary Fig. S6D). An increase in the content of several polyhydroxy FA isomers was also observed in the leaves of *gl15-S* mutant (Supplementary Fig. S6E). These results provide a further indication of the epistatic effect of the *gl15-S* recessive allele on the *ZmFDL1* gene.

Differently, both *gl15-S* and *fdl1-1* mutations caused a significant decrease in the total wax load, as compared to wild-type plants (Fig. 5E). This decrease was enhanced in the *fdl1-1 gl15-S* mutant, thus suggesting an additive effect of the two genes on wax abundance. This effect was conserved also examining the specific classes of wax metabolites except for the total alkanes load that did not statistically differ among the four genotypic classes (Fig. 5E). The total aldehydes content was reduced in the double *fdl1-1 gl15-S* mutant compared to wild type but not in single mutants (Fig. 5E). Contrariwise, the abundance of long-chain alcohols, which was diminished by both *gll15-S* and *fdl1-1* mutations, showed equal levels in *fdl1-1 gl15-S* and *gl15-S* mutants (Fig. 5E). Differences between genotypes were also detected analysing the wax relative abundance of specific class of metabolites. Compared to wild-type plants (3.4%), a progressive increase in alkane percentages was found in *fdl1-1* (4.1%), *gl15-S* (5.5%) and *fdl1-1 gl15-S* (6.1%), respectively (Supplementary Fig. S6B). The relative abundance of aldehydes was lower in the *fdl1-1* mutant (12.6%) compared to the wild-type control (17.9%) and was partially restored in the *fdl1-1 gl15-S* mutant (15.5%) but was statistically higher in *gl15-S* mutant (26.2%) (Supplementary Fig. S6B). Relative to the total wax abundance, the long-chain alcohols appeared higher in *fdl1-1* and lower in *gl15-S,* compared to wild type (78.6%), with a percentage of 83.2% and 68.3%, respectively (Supplementary Fig. S6B). These differences were lost in the *fdl1-1 gl15-S* mutant (78.3%) which was similar to the wild-type control (Supplementary Fig. S6B).

Various trends were observed for the specific wax metabolites (Supplementary Fig. S6F-H). For instance, C32 n-aldehydes (ALD32; Supplementary Fig. S6G) and C24 (24OH) and C30 (30OH) alcohols (Supplementary Fig. S6H) were lowest in the *fdl1-1 gll15-S* as compared to the other genotypes. The abundance of C30 n-aldehydes (ALD30; Supplementary Fig. S6G) was similar in *fdl1-1* single and *fdl1-1 gl15-S* mutant plants, while considering C32 alcohols, *gl15-S* single and *fdl1-1 gl15-S* double mutants revealed no statistical differences (32OH; Supplementary Fig. S6H). These results suggest that the genetic constitution of both *ZmGL15* and *ZmFDL1* genes affects the accumulation of wax metabolites and the action of the two genes is exerted in different ways on different compounds.

Overall, the analysis of the lipid profiles indicates that in maize the chemical composition of the juvenile cuticle is affected by the genetic interactions between ZmGL15 and ZmFDL1 regulatory factors.

To investigate the consequences of the differences in both loads and chemical composition on the epicuticular wax crystals density and distribution, the surface of the second leaf was examined in the four genotypic classes by SEM analysis (Fig. 6; Supplementary Fig. S7). Despite the dull appearance of the second leaf, the *gl15-S* mutant presented less wax crystals (Fig. 6J; Supplementary Fig. S7J), as we previously observed in the third glossy leaf (Fig. 3J, L). The epicuticular alterations observed on both adaxial (Fig. 6B, F, J) and abaxial (Supplementary Fig. S7B, F, J) leaf surfaces, in *gl15-S* mutants were similar to those of *fdl1-1 gl15-S* mutants (Fig. 6D, H, L; Supplementary Fig. S7D, H, L). Specifically, the surfaces of both genotypes displayed a lower density of wax crystalloids, and these crystalloids were more irregular and flattened than the wild-type control (Fig. 6A, E, I; Supplementary Fig. S7A, E, I) and the *fdl1-1* mutant (Fig. 6C, G, H; Supplementary Fig. S7C, G, H). The area of the amorphous regions appeared even more extended in the *fdl1-1 gl15-S* mutant supporting the idea of an additive effect of the *ZmFDL1* and *ZmGL15* genes on wax biosynthesis. Differently, in the *fdl1-1* mutant, no alteration in epicuticular wax crystals density and distribution was detected (Fig. 6C, G, H; Supplementary Fig. S7C, G, H) suggesting that the reduction in the total wax content (Fig. 5E) can alter the cuticle-dependent leaf permeability (Fig. 5B) but is not sufficient to induce ultrastructural alterations. Altogether, these data provide further indications that the phenotype conferred by the *gl15-S* mutation is epistatic to the fdl1 phenotype. They also show that both the cuticle-mediated leaf permeability and the epicuticular ultra-structure are influenced by the genetic interaction between the two genes.

**Figure 6.**
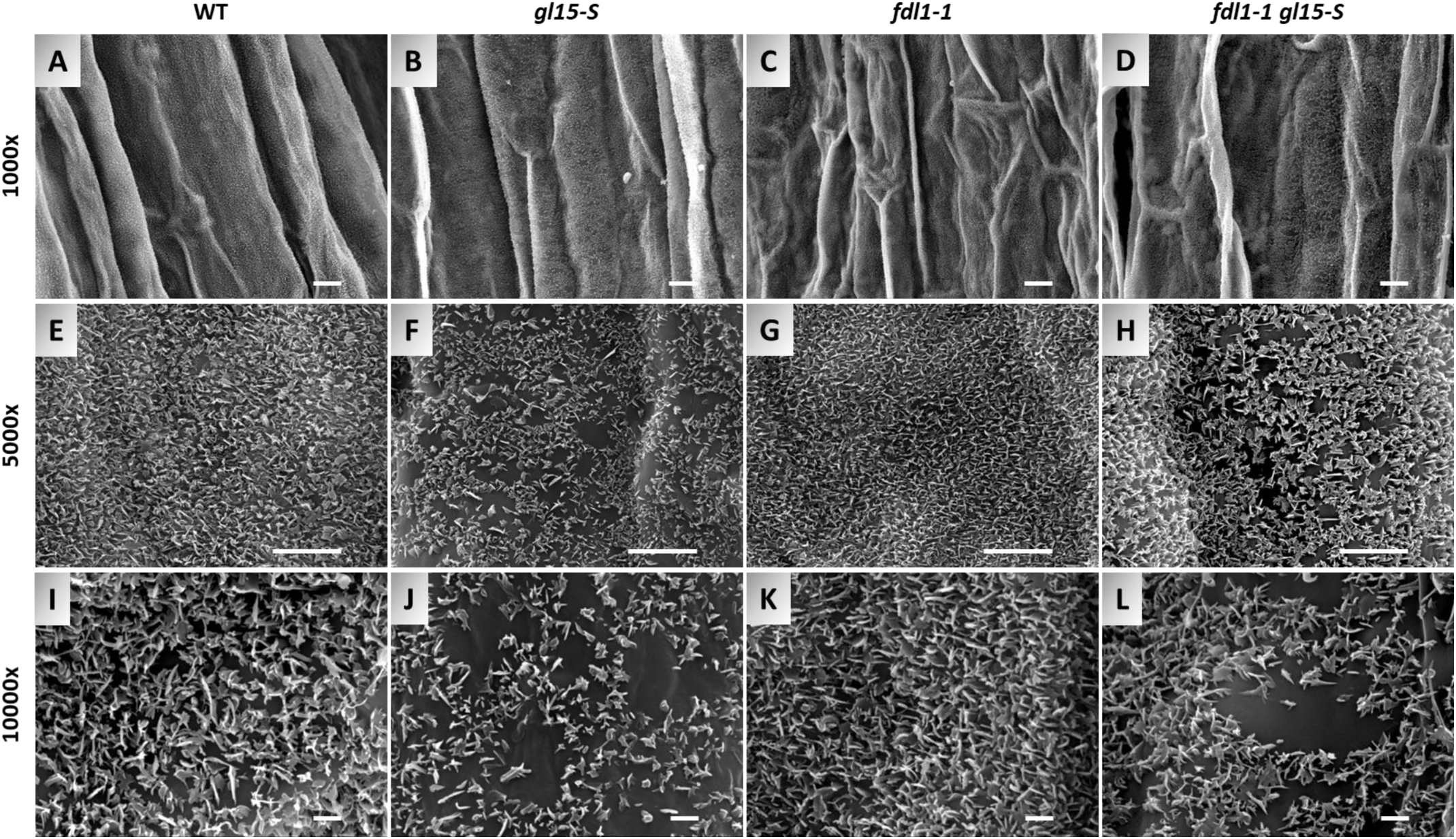
Distribution of cuticular waxes on the adaxial leaf surface of the *fdl1-1 gl15-S* double mutant. SEM micrograph images of the adaxial surface of the second fully expanded leaf in wild type (WT) control plant (A, E, I), single homozygous *gl15-S* (B, F, J), single homozygous *fdl1-1* (C, G, K) and double homozygous *fdl1-1 gl15-S* (D, H, L) mutants have been acquired at 1000x, 5000x and 10000x magnification. Scale bars correspond to 10 µm (A-D), 5 µm (E-H) and 1 µm (I-L).

## DISCUSSION

### Spatiotemporal *ZmFDL1* and *ZmGL15* transcript pattern analysis revealed co-expression in the juvenile epidermis and reciprocal stimulation

Previous studies have indicated that the *FUSED LEAVES1 (FDL1)* gene plays a key role on the control of juvenile cuticle deposition in maize (La Rocca *et al*., 2015; Castorina *et al*., 2020). The *ZmFDL1* transcript level is higher in the coleoptile and first leaf, while showing a progressive decrease in the second and third leaf. In addition, the homozygous *fdl1-1* mutant plants exhibited a defective phenotype in the first stages of growth, whereas, following the third or fourth leaves stage, they resumed a normal phenotype and were indistinguishable from wild-type siblings (La Rocca *et al*., 2015). The cuticle-related chemical defects appeared transiently in germinating mutant seedlings (up to the second leaf stage), and a progressive shift to control values was observed in subsequent plant developmental stages (Castorina *et al*., 2020). It was thus proposed that the action of ZmFDL1 was dispensable at later stages of development and replaced by that of other regulatory genes. We have also proposed that cuticle deposition during the juvenile phase is due to the interaction between ZmFDL1 and other regulatory factors involved either in maintaining the juvenile phase or in promoting the transition from juvenile to adult phase (Castorina *et al*., 2020). In this context, the *GLOSSY15 (GL15)* gene (Moose and Sisco 1994) was considered as an interesting candidate. This hypothesis was corroborated by the observation that *ZmGL15* was down-regulated in the *fdl1-1* mutant (Castorina *et al*., 2020). Moreover, *ZmGL15* was demonstrated to be crucial for the maintenance of the juvenile epidermal traits (Lauter *et al*., 2005).

*ZmFDL1* is much more highly expressed in the epidermis, where the cuticle components are synthetised, than in other tissue layers (Liu *et al*., 2020; Fig. 1B). In contrast, we found that *ZmGL15* is equally transcribed in the different leaf cell types, including the epidermal cells. Since *ZmGL15* acts in a cell-autonomous manner to specifically regulate the juvenile leaf epidermal traits (Moose and Sisco 1994), we propose that in the epidermis *ZmGL15* specifically controls the expression of different cuticle related-genes, among which *ZmFDL1* (Fig. 4C). Moreover, the observation that *ZmFDL1* was down regulated in the *gl15* mutants (Fig. 1C) reveals that in the epidermis the two genes reciprocally stimulate their expression. On this basis, we have further investigated the genetic relationship between these two transcription factors.

### The adult-like cuticle restores the wax-dependent cuticular alterations and exhibits increased leaf water-holding capacity

We first obtained evidence that *ZmGL15* plays a crucial role in modulating the properties of the juvenile cuticle. We examined the *gl15-S* first and second leaves, which exhibited a dull phenotype and juvenile epidermal traits, and the third leaf, which was in transition and showed a glossy phenotype (Fig. 2A; Supplementary Fig. S2). Therefore, the *gl15-S* mutant was used as a genetic tool to first characterize the differences between juvenile and adult cuticle in maize.

A decrease in the total wax load and several changes in wax composition (Fig. 4; Supplementary Fig. S4; Fig. 5; Supplementary Fig. S6) were observed, confirming the results obtained in a previous study conducted for a wax-related mutant, also named *gl15*, which showed an altered wax composition (Avato *et al*., 1987). Strikingly, some features of the *gl15-S* cuticle, such as a higher proportion of alkanes (Supplementary Fig. S4B, F), the reduced abundance of 32 aldehydes (Supplementary Fig. S4G) and of long-chain alcohols (Supplementary Fig. S4B, H; Supplementary Fig. S6B) make the wax composition of the *gl15-S* mutant cuticle in the analysed leaves more similar to that of an adult maize leaf (Bianchi *et al*., 1984; Avato *et al*., 1990; Yang *et al*., 1993; Bourgault *et al*., 2020). The reduction in the cuticular waxes is consistent with the reduction in size and distribution of the epicuticular wax crystals, which was observed by SEM analysis of leaf epidermis (Fig.3 and Fig.6). The contribution of *ZmGL15* to wax deposition was also confirmed by the changes in the transcript level of a group of wax-related genes in *gl15-S* mutant leaves (Fig. 4). Among these, ZmGL3 is a MYB transcription factor that affects the expression of several genes involved in the biosynthesis of very-long-chain fatty acids (Liu *et al*., 2012) and ZmGL13 is an ABC transporter required for the accumulation of epicuticular waxes (Li *et al*., 2013). We additionally analysed the cutin component and observed an increase in the cutin load in all *gl15-S* mutant leaves examined (Fig. 4A; Fig. 5D) that was due to an increase in ω-hydroxy fatty acids (FAs) as well as in polyhydroxy FAs (Fig. 4A; Fig. 5D).

Similar to *gl15-S,* the *fdl1-1* mutation caused a significant decrease in the total wax load in the second leaf, as compared to wild-type plants (Fig. 5D), due to decreased aldehydes and long-chain alcohols (Fig. 5E). Nevertheless, unlike *gl15-S*, the *fdl1-1* mutation did not affect the total cutin load in mature leaf as compared to that of wild-type siblings (Fig. 5D). The slight decrease observed for FAs was perhaps balanced by a small increase of both ω-hydroxy FAs and polyhydroxy FAs (Fig. 5D).

The greater reduction in the total wax load observed in the *fdl1-1 gl15-S* double mutants (Fig. 5E), suggests that the two genes have an additive effect on this trait. In addition, a partial epistatic effect of *gl15-S* on *fdl1-1* was observed for the cutin load, since in the double mutants it was higher than in homozygous *fdl1-1.* Based on these data, a model has been proposed to explain how the genetic interaction between *ZmFDL1* and *ZmGL15* modulates the deposition of the juvenile cuticle. The formation of a juvenile cuticle is positively stimulated by the action of both *ZmFDL1* and *ZmGL15* (Fig. 7B, top left), which promotes waxes biosynthesis. Furthermore, the action of ZmGL15 in maintaining the juvenile vegetative phase generates a temporal window of competence that ensures a high level of *ZmFDL1* expression, thus allowing it to fully express its role in promoting a juvenile cuticle. Consequentially, the action of *ZmGL15* prevails on that of *ZmFDL1* in the establishment of the juvenile cuticle (Fig. 7C).

**Figure 7.**
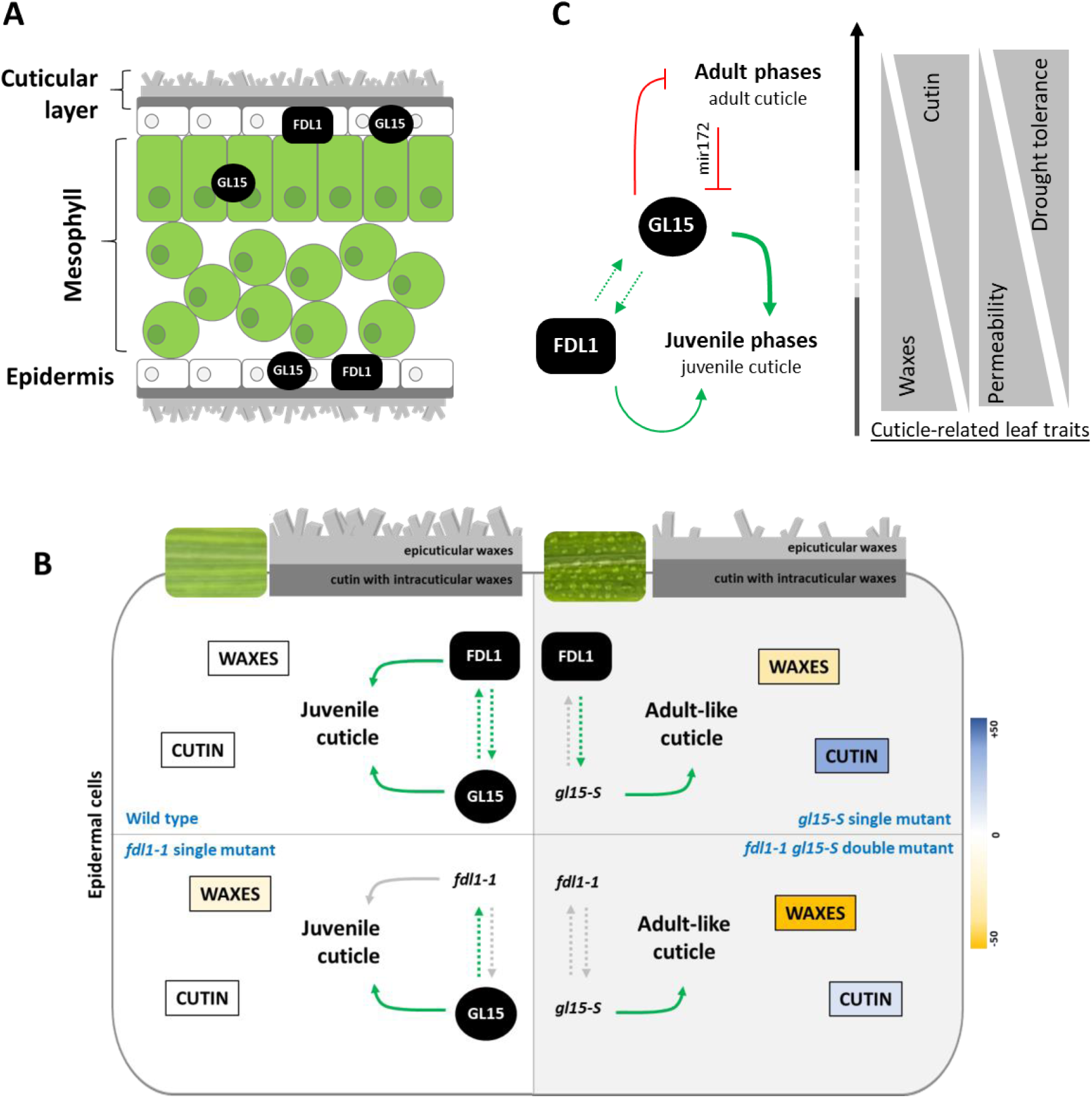
Proposed roles of ZmFDL1 and ZmGL15 transcription factors in the regulation of cuticle deposition in the juvenile leaves. (A) *ZmGL15* is expressed, during the juvenile vegetative phase, in both the epidermal and the mesophyll cells to maintains the juvenile leaf traits. The transcript of *ZmFDL1* is specifically expressed in the epidermis and regulates the cuticle deposition in the juvenile leaves. (B) Genetic-dependent *ZmFDL1* and *ZmGL15* interaction leads to proper modulation of juvenile cuticle relative composition. The cuticle of wild-type plants is rich in waxes which confer the characteristic dull phenotype to the juvenile leaves. Conversely, the adult-like cuticle of the *gl15-S* mutants is rich in cutin and poor in waxes, giving the leaves a glossy appearance. (C) ZmFDL1 and ZmGL15 integrate leaf developmental stimuli in the cuticle biosynthetic pathway through the reciprocal activation of their expression, thus ensuring the correct biosynthesis and deposition of cuticular compounds peculiar to the juvenile cuticle. The total wax content decreases in adult leaves compared to juvenile leaves, while the amount of cutin increases to reduce cuticle-dependent leaf permeability. A cuticle enriched in cutin reduces the non-stomatal leaf evapotranspiration and prevents water loss, thus improving the tolerance to drought. Linear green arrows indicate transcriptional activation and red T-shaped lines indicate transcriptional repression. Curved green arrows indicate positive regulation of the cuticle pathways. Colour scale reflects the magnitude of total level for each class of compounds. Yellow and blue colors indicate reduced and increased amounts of cuticular compounds respect to wild type, respectively.

In the absence of the *ZmGL15* action, despite the genetic contribution of *ZmFDL1,* the cuticle properties are more similar to that of an adult-like cuticle, with lower wax and higher cutin content (Fig. 7B top right, C). Despite the aforementioned alterations, a reduction in the chlorophyll leaching rate was observed in the *gl15-S* mutant, which is indicative of a less permeable cuticle (Fig. 2B; Supplementary Fig. S3 A, C; Fig. 5B, C). More interestingly, under drought stress, the leaf RWC was higher in leaves of the *gl15-S* mutant as compared to wild-type control siblings (Fig. 2D; Supplementary Fig. S3G). This parameter, which reflects the balance between water supply to the leaf tissue and transpiration rate, can be considered as an indicator of the water-holding capacity and water status in plants (Lugojan and Ciulca 2011). It is therefore conceivable that a the less permeable cuticle caused by the *gl15-S* mutation contributes to increasing the water-holding capacity of the leaf.

In the absence of the *ZmFDL1* action, the juvenile cuticle composition is affected with a decrease in the total wax load (Fig 5D, E; Fig. 7B bottom right, C). The aforementioned alterations resulted in a higher chlorophyll leaching, which indicates that the *fdl1-1* mutant has a more permeable cuticle (Fig. 5B, C). The *fdl1-1* defective cuticle-dependent leaf permeability was rescued in the double-homozygous *fdl1-1 gl15-S* mutant in which the chlorophyll leaching rate resulted lower than wild type and equivalent to that of the single-homozygous *gl15-S* mutant (Fig. 5B, C). Despite the significant reduction in the total wax load observed in the double mutant, the *gl15*-*S* dependent high enrichment in cutin is sufficient to overcome the wax alterations and renders the cuticle less permeable (Fig. 7B bottom right, C).

### Cutin is a key component in the constitution of an effective cuticular water barrier in leaf maize seedlings

The cuticle, composed of cutin and cuticular waxes, provides structural integrity to the plant surface. A well-formed cuticle is known to help maintaining cell turgor and preventing desiccation under drought conditions (Kerstiens, 2006). However, the complex relationships between cuticle composition, structure, and water barrier function are still poorly understood.

Most of the studies have considered the role of cuticular wax amount in the response to drought (Shepherd and Wynne Griffiths 2006; Cameron *et al*., 2006; Bourdenx *et al*., 2011; Zhou *et al*., 2013; Zhu and Xiong 2013; Lee and Suh, 2022; Li *et al*., 2019) and few or none the role of the cutin fraction. Often, the cuticular waxes are identified as the actual barrier of plant cuticle against the diffusion of water and the cutin matrix is considered as a kind of framework within which waxes are deposited and crystallized. It has been proposed that mutations that destroy the cutin framework potentially disrupt wax crystallization, and subsequently the permeability function of the cuticle. However, the dehydration-avoidant ZPBL 1304 maize line exhibited significantly lower rates of leaf water loss and thicker cuticle compared to the dehydration-susceptible ZPL 389 maize line, but contrary to expectations it had less cuticular waxes (Ristic and Jenks, 2002). These results indicated that that water flow through the cuticle is complex and that the amount of cuticular wax alone does not determine the rate of epidermal water loss, suggesting that other factors are involved (Ristic and Jenks, 2002).

The main role of cutin was first confined to plant development where it is responsible for correct organs separation. This was evidenced by the studies of cutin defective mutants which frequently display organ fusions (Bird *et al*., 2007; Ingram *et al*., 2017). This role was also confirmed by our data: the *gl15*-dependent increase in the total cutin load have significantly alleviated the developmental defects observed in the *fdl1-1* seedlings (Supplementary Fig. S5C-D). Nevertheless, it is still largely unclear what role cutin plays in cuticle-dependent leaf permeability to restrict transpirational water loss.

Cutin is the most abundant structural component of the cuticle (Pollard *et al*., 2008; Beisson *et al*., 2012). Several transcription factors have been identified (Kannangara *et al*., 2007, Yeats and Rose, 2013; Suh *et al*., 2022) as playing a crucial role in the regulation of the expression of cutin biosynthesis genes (Yeats *et al*., 2012; Girard *et al*., 2012; Yeats *et al*., 2014; Philippe *et al*., 2020). In tomato, the fruits of the *cutin deficient 1* (*cd1*) mutant showed a significantly increased transpiration rate (Isaacson *et al*., 2009) and the leaf of the *sticky peel* (*pe*) mutant exhibited increased cuticular permeability (Nadakuduti *et al*., 2012). In Arabidopsis, the loss of cutin monomers in *gpat4*/*gpat8* mutant plants resulted in an increase in cuticle permeability in the presence of normal wax loads (Li *et al*., 2007) and this was consistent also with previous observations on the *att1* and *lacs2* cutin mutants (Schnurr *et al*., 2004; Xiao *et al*., 2004). More recent evidence has highlighted that, under drought stress, the expression of genes involved in cutin biosynthesis are often upregulated and a functional cutin is essential to cope with environmental stresses (Chen *et al*., 2011; Fich *et al*., 2016). The importance of cutin for the water barrier function is also substantiated by the comparison of the cuticle composition in maize adult leaves at various developmental stages, which showed that cutin amount correlates with cuticle-dependent leaf transpiration. The reduction in cuticle permeability during leaf development coincides with an increase in the deposition of the cutin polyester, thus implying that the cutin-rich layer is a key component to establish the water barrier property of mature cuticle (Bourgault *et al*., 2020).

Moreover, cuticle amount is modulated in response to changes in the environment. In Arabidopsis, water deficit-treated plants showed 75% more wax than control plants and also a 65% increase in total cutin (Kosma *et al*., 2009). In general, alkanes accounted for the observed large increases in total wax amount, whereas nearly all cutin monomers increased. This indicates that the total cutin load, rather than the amount of any specific cutin constituent, is of importance in the water deficit stress response (Kosma *et al*., 2009).

In our study, we showed that a cutin-enriched cuticular layer (Fig. 4A, 5D) combined with the accumulation of specific wax types, such as alkanes (Fig. 4B, 5E), improves the properties of the cuticle (Fig. 2B, 5B), thus forming an effective hydrophobic barrier that limits water loss, thereby helping the plant in retaining moisture (Fig. 2D). A cuticular layer with a greater cutin load is more efficient in preventing leaf water loss. Overall, the results demonstrate that the cuticle composition, more than its total amount, is the key factor in determining the non-stomatal leaf transpiration. Moreover, our data indicate that an improved cuticle, with an increased amount of cutin, is also able to compensate the leaf permeability alterations due to severe wax reduction. On this basis, we propose that cutin represents a crucial component of the cuticle in establishing an effective water barrier layer, and consequently has a positive effect on plant adaptation to water scarcity conditions. These findings provide insights into the complex relationships between cuticle composition, structure, and function in maize leaves, highlighting the importance of specific lipid components in establishing the water barrier property. They corroborate a first idea, which refuted that cuticle permeability depends only on wax load or cuticle thickness, and proposed that cuticle composition and the organization of its components determine water barrier function (Riederer and Schreiber 2001).

There is an increasing interest in breeding for cuticle-related traits that promote drought tolerance. Improving cuticle can reduce water loss and protect plants from desiccation, which is crucial for maintaining crop productivity in the context of climate change. Drought tolerance in crops like wheat, maize, and rice, which are staple foods for a large portion of the global population, can lead to more stable yields also in regions prone to water scarcity (Kan *et al*., 2022; Busta *et al*., 2021). The insights gained from this research can drive the selection of desirable traits in crop breeding, speeding up the development of stress-resistant varieties.

### Supplementary data

The following Supplementary data are available at JXB online. Supplementary Figures S1-S7 and Table S1 are collected in a pdf supplementary file: Table S1. Primers used for quantitative Real-Time PCR analysis; Figure S1. Expression pattern of *ZmFDL1* and other cuticle genes; Figure S2. Epidermal traits in *gl15-S* and wild-type leaves; Figure S3. Cuticle-dependent leaf permeability on homozygous *gl15-S* and wild-type control plants; Figure S4. Detailed cuticle composition in first, second and third leaves of *gl15-S* mutant; Figure S5. Epistatic interaction between *ZmGL15* and *ZmFDL1*; Figure S6. Detailed cuticle composition in the second fully expanded leaf from 15-day-old plants; Figure S7. Distribution of cuticular waxes on the abaxial leaf surface of the *fdl1-1 gl15-S* double mutant.

## Acknowledgements

The authors express sincere gratitude to Stephen P. Moose for generously providing the *gl15* maize mutant lines used in this study. The contribution of these valuable genetic resources has been instrumental in advancing our research. The authors also thank Stefania Crespi, Department of Earth Sciences “Ardito Desio”-Università degli Studi di Milano for the SEM images acquisition and the Bordeaux□Metabolome Facility, which is supported by MetaboHUB (ANR□11□INBS□0010), where all fatty acyl GC-based analyses were performed.

## Author contributions

CaG and CoG: conceptualization; CaG and DF: methodology; CaG: formal analysis; CaG and DF: investigation; CoG and FD: resources; CaG and DF: data curation; CaG: visualization; CaG: supervision; CaG and CoG: writing - original draft; All: writing - review & editing; All authors read and approve the final manuscript.

## Conflict of interest

The authors declare no conflict of interest.

## Funding

This study was carried out within the Agritech National Research Center and received funding from the European Union Next-GenerationEU (PIANO NAZIONALE DI RIPRESA E RESILIENZA (PNRR) – MISSIONE 4 COMPONENTE 2, INVESTIMENTO 1.4 – D.D. 1032 17/06/2022, CN00000022). This manuscript reflects only the authors’ views and opinions, neither the European Union nor the European Commission can be considered responsible for them.

## Data availability

All data are available under request to the authors.

## Tables

**Supplementary Table 1. Primers used for quantitative Real-Time PCR analysis.** The gene models are referred to the Zm-B73-REFERENCE-NAM-5.0 reference genome version. The asterisk on the gene symbol indicate the gene use as housekeeping to normalize the gene expression in RT-qPCR assay.

**Supplementary Figure 1. Expression pattern of *ZmFDL1* and other cuticle genes.** Epidermis enrichment test of cuticle-related gene expression. Transcript level analysis of *ZmCER1, ZmCER4, ZmONI3, ZmHTH1, ZmWSD11, ZmKCS16, ZmKCS39, ZmGL1, ZmGL2, ZmGL3, ZmGL4, ZmGL8, ZmGL13* and *ZmGL14* genes was performed, by RT-qPCR, on the whole leaf (Total) and on the manually dissected (by peeling) epidermal tissue from the second leaves of 10-day-old wild-type plants. Values represent the mean fold change of a minimum of four biological replicates. Error bars are ±SE. Comparison was made between genotypes and significant differences were assessed by Student’s T-test (* P<0.05; ** P<0.01; *** P<0.001; **** P<0.0001; ns=not significant).

**Supplementary Figure 2. Epidermal traits in *gl15-S* and wild-type leaves.** Glue-imprinted leaf surface (A, C, E, G, I, K, M, O) and Toluidine blue-stained epidermal peels (B, D F, H, J, L, N, P) from the adaxial and abaxial side of the second and third fully expanded leaves in wild-type (WT) (A-D, I-L) and homozygous *gl15-S* mutant (E-H, M-P) plants. Juvenile epidermal cells uniformly stain violet/pink, and possess wavy cell walls (K, inset). Epidermal cells with adult traits show aqua/turquoise-staining cells with highly crenulated lateral walls (O, inset). Transition leaves present epidermal cells with both juvenile and adult characteristics. Images were acquired with light microscopy. Scale bars correspond to 100 µm (A, C, E, G, I, K, M, O) and 50 µm (B, D F, H, J, L, N, P).

**Supplementary Figure 3. Cuticle-dependent leaf permeability on homozygous *gl15-S* and wild-type control plants.** The chlorophyll leaching assay was performed on the first (A), third (C) and fourth (E) fully expanded leaves of homozygous *gl15-S* and wild-type (WT) control plants. Leaf evapotranspiration in the first (B), third (D) and fourth (F) detached leaves is expressed as the percentage of water loss of fresh weight (FW). Values represent the mean ± SE of 10 biological replicates per genotype. Significant differences were assessed by Student’s T-test (\**P*<0.05; \*\**P*< 0.01; \*\*\**P*<0.001; and \*\*\*\**P*<0.0001; ns, not significant). (G) The relative water content (RWC) was measured in the third leaf of wild-type (WT) and *gl15-S* plants subjected to 24 and 48 hours of water scarcity imposed by withholding irrigation. Values represent the mean of a minimum of four biological replicates. Error bars are ±SE. Different letters denote significant differences assessed by two-way ANOVA (*P*<0.05). (H) Stomatal index and (I) the density per 1 mm^2^ of guard cells (GC) and pavement cells (PC) were measured in both the abaxial and adaxial sides of the third leaf of wild-type (WT) and *gl15-S* plants. Values are the mean ± SE and differences were evaluated by Student’s T-test.

**Supplementary Figure 4. Detailed cuticle composition in first, second and third leaves of *gl15-S* mutant.** Percentage of (A) cutin and (B) wax compound classes relative to the total abundance. Black, white and grey letters above or within the data bars indicate statistically significant differences between genotypes as assessed by two-way ANOVA (*P*<0.05) for comparisons within a specific class of cuticular compound. Relative amounts of (C) fatty acids (FAs), (D) ω-hydroxy fatty acids, (E) polyhydroxy-fatty acids, (F) alkanes, (G) aldehydes and (H) long-chain primary alcohols. Significant differences were assessed by Student’s T-test (* *P*<0.05, ** *P*<0.01, *** *P*<0.001, **** *P*<0.0001, ns=not significant). Values represent the mean of independent biological replicates (N=6) for wild-type (WT) and single homozygous *gl15-S* mutant plants. Error bars are ±SE.

**Supplementary Figure 5. Epistatic interaction between *ZmGL15* and *ZmFDL1*.** (A) Representative phenotypes of 15-day-old wild-type (WT) control plant, *gl15-S*, *fdl1-1* and *fdl1-1 gl15-S* mutant seedlings. (B) Representative phenotype of 15-day-old wild-type (WT) and *fdl1-1* seedlings. Numbers 1, 2, 3 and 4 represent mild, intermediate, strong and severe fdl1 phenotypes, respectively. (C-D) The pie charts reported the frequency of each phenotypic class present in (C) single homozygous *fdl1-1* and (D) double homozygous *fdl1-1 gl15-S* mutant. (E) Box plot of the flowering time expressed as total leaf number measured at maturity. Values represent the mean of independent biological replicates per genotype (minimum N=9). Different letters denote significant differences between genotypes assessed by two-way ANOVA (*P*<0.05). (F-Q) Toluidine blue-stained epidermal peels dissected from the abaxial side of the second (F-I), third (J-M) and fourth (N-Q) fully expanded leaves in wild-type (F, J, N), single homozygous *fdl1-1* (G, K, O) and *gl15-S* mutant (H, L, P), and double homozygous *fdl1-1 gl15-S* mutant (I, M, Q) plants. Scale bars correspond to 50 µm.

**Supplementary Figure 6. Detailed cuticle composition in the second fully expanded leaf from 15-day-old plants.** Percentage of (A) cutin and (B) wax compound classes relative to the total abundance. Relative amounts of (C) very-long-chain fatty acids (VLCFAs), (D) ω-hydroxy fatty acids, (E) polyhydroxy-fatty acids, (F) alkanes, (G) aldehydes and (H) long-chain primary alcohols. Values represent the mean of independent biological replicates (N=6) for wild-type (WT), single homozygous *fdl1-1* and *gl15-S,* and double homozygous *fdl1-1 gl15-S* mutant plants. Error bars are ±SE. Different letters above data bars denote statistically significant differences between genotypes assessed by two-way ANOVA (*P*<0.05). In panels A and B, black, white and grey letters above or within the data bars represent comparisons within a given class of cuticular compound.

**Supplementary Figure 7. Distribution of cuticular waxes on the abaxial leaf surface of the *fdl1-1 gl15-S* double mutant.** SEM micrograph images of the abaxial surface of the second fully expanded leaf in wild type (WT) control plant (A, E, I), single homozygous *gl15-S* (B, F, J), single homozygous *fdl1-1* (C, G, K) and double homozygous *fdl1-1 gl15-S* (D, H, L) mutants have been acquired at 1000x, 5000x and 10000x magnification. Scale bars correspond to 10 µm (A-D), 5 µm (E-H) and 1 µm (I-L).

## Notes

### Competing Interest Statement

The authors have declared no competing interest.

